# OMA standalone: orthology inference among public and custom genomes and transcriptomes

**DOI:** 10.1101/397752

**Authors:** Adrian M Altenhoff, Jeremy Levy, Magdalena Zarowiecki, Bartłomiej Tomiczek, Alex Warwick Vesztrocy, Daniel A Dalquen, Steven Müller, Maximilian J Telford, Natasha M Glover, Christophe Dessimoz

## Abstract

Genomes and transcriptomes are now typically sequenced by individual labs, but analysing them often remains challenging. One essential step in many analyses lies in identifying orthologs—corresponding genes across multiple species—but this is far from trivial. The OMA (Orthologous MAtrix) database is a leading resource for identifying orthologs among publicly available, complete genomes. Here, we describe the OMA pipeline available as a standalone program for Linux and Mac. When run on a cluster, it has native support for the LSF, SGE, PBS Pro, and Slurm job schedulers and can scale up to thousands of parallel processes. Another key feature of OMA standalone is that users can combine their own data with existing public data by exporting genomes and pre-computed alignments from the OMA database, which currently contains over 2100 complete genomes. We compare OMA standalone to other methods in the context of phylogenetic tree inference, by inferring a phylogeny of the Lophotrochozoa, a challenging clade within the Protostomes. We also discuss other potential applications of OMA standalone, including identifying gene families having undergone duplications/losses in specific clades, and identifying potential drug targets in non-model organisms. OMA Standalone is available at http://omabrowser.org/standalone under the permissible open source Mozilla Public License Version 2.0.

## Introduction

The sequencing revolution is yielding a flood of genomes and transcriptomes, with thousands already sequenced and many more underway (Pagani et al., 2012). A powerful way of characterising newly sequenced genes is to compare them with evolutionarily related genes—in particular with orthologs in other species (Dessimoz et al., 2012; Forslund et al., 2017; Sonnhammer et al., 2014). In this way, experimental knowledge from model organisms can be propagated to non-model organisms. Elucidation of orthology and paralogy relationships is also essential to reconstruct species trees, to better understand the mechanics of gene/genome evolution, to study adaptation, or to pinpoint the emergence of new gene functions (Gabaldón and Koonin, 2013).

The importance of determining orthology has led to the development of many inference methods and associated databases (reviewed in Altenhoff and Dessimoz, 2012). Some of the best established orthology resources include EggNOG (Huerta-Cepas et al., 2016), Ensembl Compara (Zerbino et al., 2018), Inparanoid (Sonnhammer and Östlund, 2015), MBGD (Uchiyama et al., 2012), OrthoDB (Zdobnov et al., 2017), OrthoMCL (Chen et al., 2006), Panther (Mi et al., 2017), PhylomeDB (Huerta-Cepas et al., 2014), and OMA (Altenhoff et al., 2017).

Key distinctive features of OMA are the high specificity of its inference pipeline (Afrasiabi et al., 2013; Altenhoff and Dessimoz, 2009; Boeckmann et al., 2011; Linard et al., 2011), the feature-rich web and programmatic interfaces, large size and taxonomic breadth of its precomputed data (currently 2167 genomes), its regular update schedule of 2 releases per year, and its sustained development over the last 13 years. The algorithms underlying the OMA pipeline have been described and validated in multiple publications (Altenhoff et al., 2013; Dessimoz et al., 2006, 2005; Roth et al., 2008; Train et al., 2017). The quality of OMA is corroborated by a recent community experiment, which highlighted the high specificity of orthologs predicted by the OMA pipeline (Altenhoff et al., 2016).

With genome and transcriptome sequencing rapidly becoming a commodity, there is an increasing need to analyse custom user data. Here, we present OMA standalone, an open-access software implementation of the OMA pipeline for Linux and Mac. We first outline some of the key features of OMA standalone. In the second part, we demonstrate the usefulness of OMA standalone in the context of species tree inference, by comparing its performance with state-of-the-art alternatives on the challenging Lophotrochozoa phylogeny.

## Results

We first highlight the defining features of OMA standalone, then turn to the phylogeny of the Lophotrochozoa, which we infer from orthologs inferred by OMA in comparison with alternative methods.

### OMA standalone software

OMA standalone takes as input the coding sequences of genomes or transcriptomes, in fasta format. The recommended input type is amino-acid sequences, but OMA also supports nucleotide sequences. With amino-acid sequences, users can combine their own data with publicly available genomes from the OMA database, including precomputed all-against-all comparisons, using the export function on the OMA website (http://omabrowser.org/export).

OMA standalone produces several types of output (also summarised in Fig. 1):

1. **Pairwise orthologs** and their subtypes (1:1, 1:many, many:1, many:many orthology). These orthologs are useful when comparing pairs of species at a time, or to identify orthologs to specific genes of interest
2. **OMA groups.** These are sets of genes for which all pairs are inferred to be orthologous. These groups are inferred as cliques (fully connected subgraphs) of pairwise orthologs. These groups are not necessarily one-to-one orthologs, but being inferred without assuming a species tree, they are particularly useful to identify marker genes for phylogenetic reconstruction.
3. **Hierarchical orthologous groups (HOGs).** These groups are defined for every internal node of the (rooted) species tree; each HOG contains the genes that are inferred to have descended from a common ancestral gene among the species attached to that internal node. Consider for instance gene ADH1, which duplicated within the primates (Carrigan et al., 2012): At the level of the last primate common ancestor, all genes that have descended from the ancestral ADH1 belong to the same HOG. However, at the level of the common ancestor of all the great apes, because ADH1 had at this point already duplicated into ADH1a, ADH1b, and ADH1c, these ancestral genes define 3 HOGs. The HOGs are stored in the standard OrthoXML format (Schmitt et al., 2011).
4. **Gene Ontology annotations.** OMA standalone annotates the input sequences with Gene Ontology annotations by propagating high-quality annotations across orthologs (Altenhoff et al., 2015). The annotations are provided in the standard GO Annotation File Format 2.1 (http://geneontology.org/page/go-annotation-file-format-20).
5. **Phylogenetic profiling.** Orthology is also used to build phylogenetic profiling—patterns of presence and absence of genes across species (Pellegrini et al., 1999). We provide two forms of output: a binary matrix with species as rows and OMA groups as columns, indicating patterns of presence or absence of genes in each group; a count matrix with species as columns and HOGs as rows, indicating the number of genes in each HOG.

**Fig. 1.**
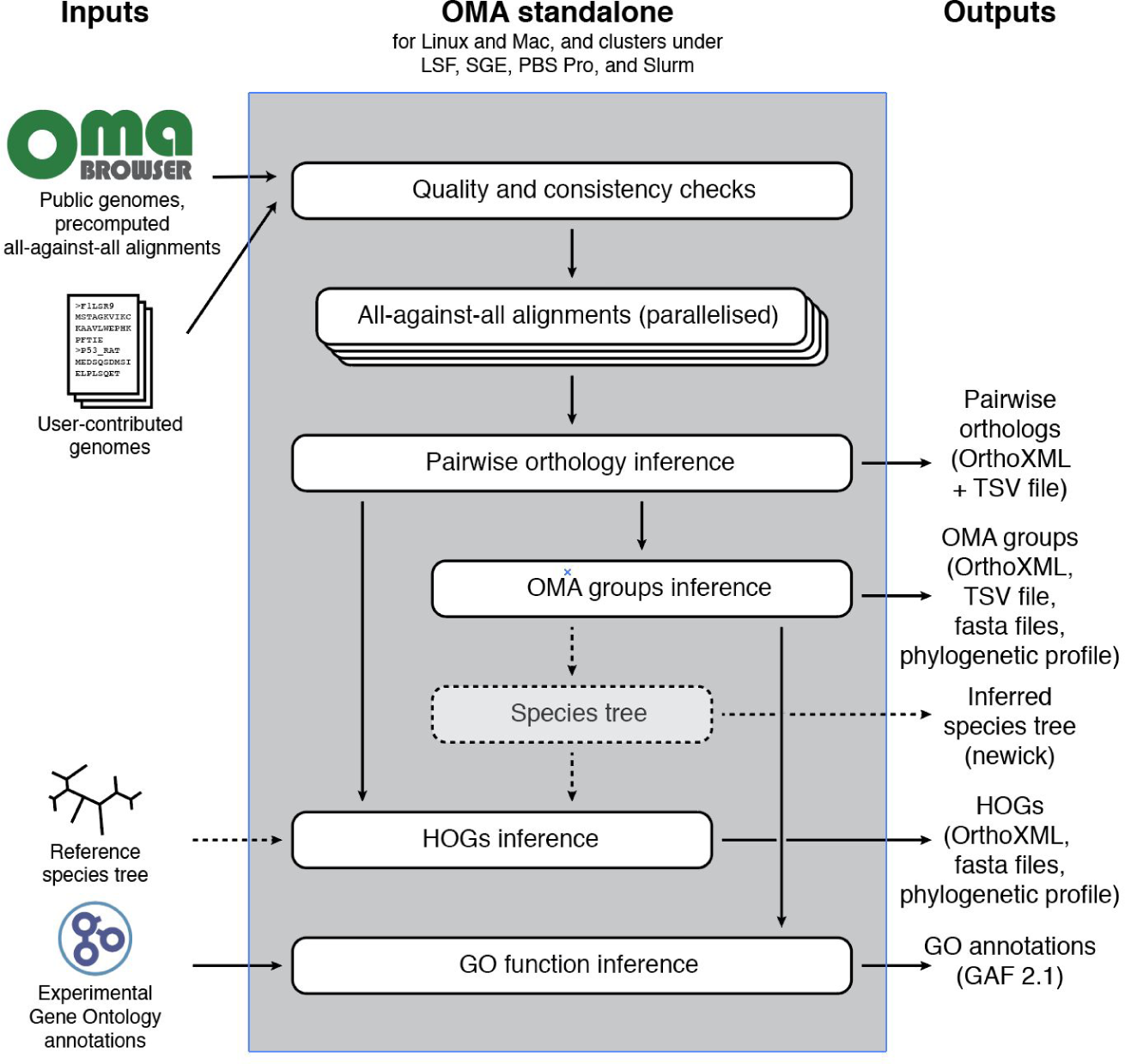
Conceptual overview of the OMA standalone software. Dotted arrows indicate alternative steps (reference species tree either specified as input or inferred from the data).

OMA standalone supports parallel computation of the all-against-all sequence comparison phase. This phase, which computes Smith-Waterman (1981) alignments followed by pairwise maximum likelihood distance estimation for all significant pairs (Roth et al., 2008), is by far the most time-consuming step of the algorithm. To fully exploit parallelism, alignments are performed using single instruction multiple data (SIMD) instructions (Szalkowski et al., 2008) on multiple cores. OMA standalone natively supports common cluster schedulers—LSF, SGE, PBS, and Slurm—and has been successfully run with several thousand jobs in parallel. Figure 2 shows typical runtimes and memory usage for datasets of various sizes.

**Fig 2:**
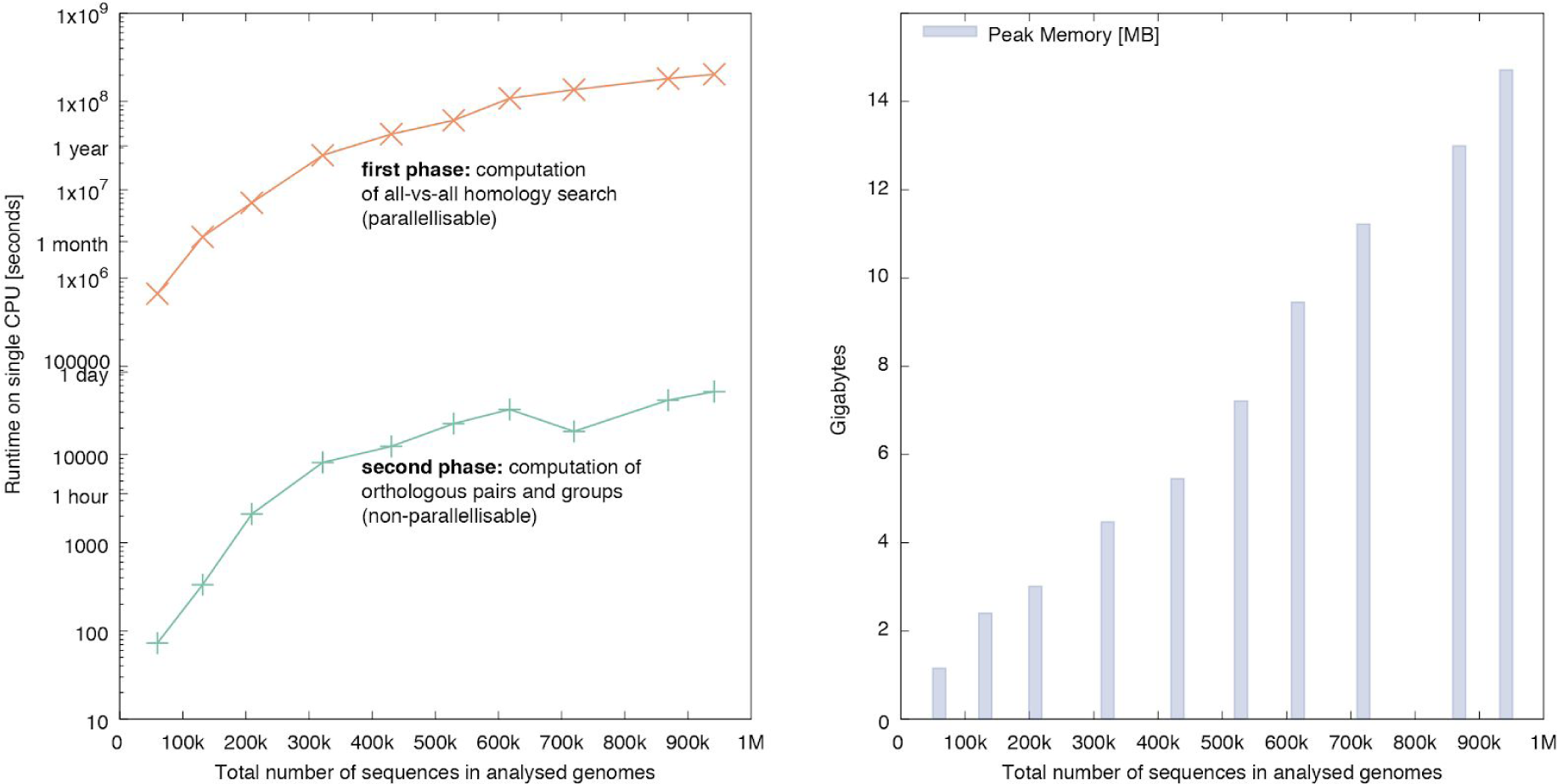
Resource measurements for various datasets of increasing sizes as total number of protein sequences. The datasets have been sampled from the public OMA Browser to maintain a constant composition of 20% fungi, 10% archaea, 10% plants, 20% metazoa and 40% bacteria genomes. **Left**: Runtime of the all-against-all phase (orange) on a single CPU, and the inference of the orthologous pairs and various groups (green). **Right**: Peak memory usage of OMA standalone in gigabytes.

### Application: the phylogenetic relationships within the Lophotrochozoa

Resolving the relationships of ancient lineages is a major challenge for molecular phylogenetics. Although some aspects of the phylogeny of the major animal clades are well resolved, the relative positions of the deeper lying clades are often disputed. The construction of large phylogenomic supermatrices, has been the method of choice for resolving the deepest nodes in the tree of life (Dunn et al., 2008; Egger et al., 2015; Fernández et al., 2014; Hejnol et al., 2009).

Fundamental to the analyses of phylogenetic relationships is the use of sequences which have descended from a single common gene in their last common ancestor, that is, orthologous sequences. Ensuring that we correctly infer orthologs is therefore vital if we are to reconstruct difficult to resolve phylogenies. The limitations of automated orthology and paralogy prediction methods with regards to phylogenetic analysis have previously been highlighted (Philippe et al., 2011b); simplistic orthology inference methods may miss orthologs (Dalquen and Dessimoz, 2013) or erroneously identify as orthologs, paralogous pairs of genes that result from differential gene losses (Dessimoz et al., 2006).

One notoriously difficult to resolve phylogeny is that of the Lophotrochozoa (Kocot, 2016), a clade of animals positioned sister to the Ecdysozoa, within the protostomes. The Lophotrochozoa contains about ten different phyla, each of which is clearly monophyletic, but the relationships between these phyla are far from clear, with many different topologies having been supported by different analyses.The inference is that the phyla are likely to have emerged in an ancient and rapid radiation resulting in weak phylogenetic signal for interphylum relationships. These circumstances make the solving of this problem particularly difficult and mean the use of accurately identified orthologs is particularly significant.

We used OMA standalone to identify orthologous marker genes among the proteomes of 19 lophotrochozoans and, as outgroups, 4 deuterostomes, 4 ecdysozoans, and 3 non-bilaterians (see Material and Methods). As a basis of comparison, we also repeated the analysis using orthology inference pipelines based on OrthoMCL (Li et al., 2003), BUSCO (Simão et al., 2015), and HaMStR (Ebersberger et al., 2009). Like OMA, these methods do not require prior specification of a species tree, are available as standalone programs and have all been used in phylogenetic analyses previously. Species trees were then constructed using these orthologs with both maximum likelihood and Bayesian tree reconstruction packages, RAxML (Stamatakis, 2014) and PhyloBayes (Lartillot et al., 2013), on the resultant supermatrices.

We first consider the amount of orthology information recovered by the various methods. OMA inferred 2,162 orthologous groups containing 15 or more species (Figure 3a). By comparison, HaMStR pipelines inferred 1,192 orthologous groups, the OrthoMCL pipeline inferred 484 orthologous groups, and BUSCO inferred 384 orthologous groups. Although OMA overall identifies more orthologous genes than other methods, it infers fewer larger groups than HaMStR and OrthoMCL. The OMA algorithm is known for having higher precision but lower recall than most other methods (Altenhoff et al., 2016). Still, in terms of total number of characters in supermatrices, OMA standalone yields a larger matrix (i.e. alignment columns) than the other methods (Figure 3b).

**Fig 3:**
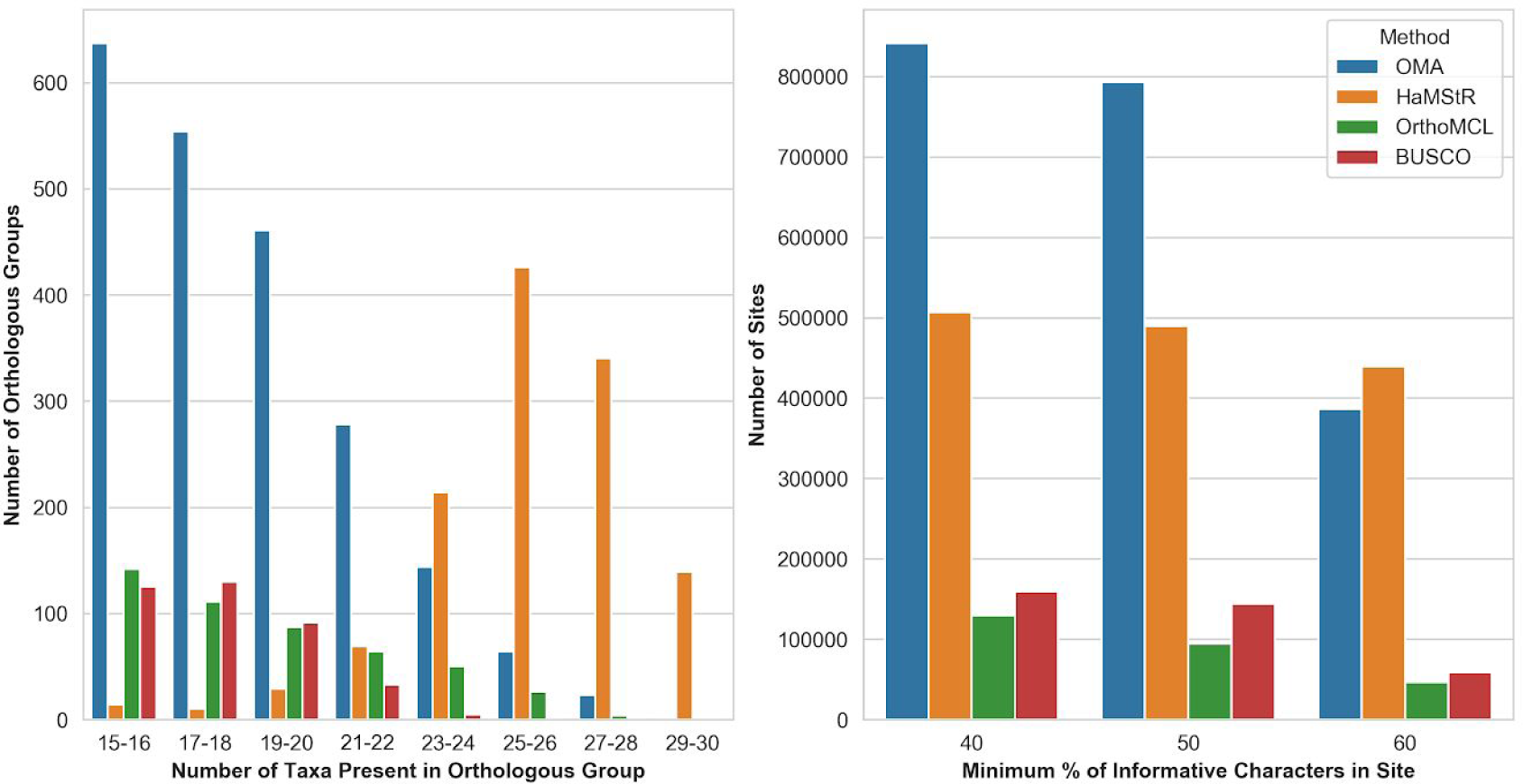
Comparison of amount of orthologous data inferred by the different pipelines. A: OMA infers more orthologous groups than other methods; the groups inferred by HaMStR are considerably larger on average than for the other methods. B: The resulting supermatrix (concatenated alignment over all orthologous groups) has most sites for OMA whether the minimum site occupancy threshold is 40% or 50%.

Using the aligned sets of orthologs identified in the previous step, we reconstructed species trees using Maximum Likelihood (RaXML, LG+I model) and Bayesian analysis (PhyloBayes, CAT+GTR+G4) on supermatrices which had been filtered to include only alignment columns with at least 60% site occupancy.

With OMA, both the RAxML tree and the Phylobayes tree had high branch support values. The RAxML tree had bootstrap support of 100 for each branch, except for five. The Deuterostomes were recovered with bootstrap support of 89, whilst the Lophotrochozoa, with the exception of Rotifera, were recovered with bootstrap support of 92. Similarly, the PhyloBayes tree had branch posterior probabilities of 1 across the tree apart from the Lophotrochozoa clade, with a posterior probability of 0.82.

The tree inferred using the ML inference method found that the Rotifera (Adineta ricciae, Brachionus plicatilis) are grouped with the Nematoda (Caenorhabditis elegans, Pristionchus pacificus), as part of the ecdysozoans. This is in disagreement with the current consensus (Giribet and Edgecombe, 2017). By contrast, the tree constructed using Bayesian inference found the Rotifera to be sister to the rest of the Lophotrochozoa, in agreement to recent studies (Egger et al., 2015; Philippe et al., 2011a). The discrepancy in the ML tree is likely due to the long branched Rotifera being attracted to the long branched Nematoda—a problem to which PhyloBayes under the CAT model has been previously shown to be more robust (Lartillot et al., 2013).

Both the ML and Bayesian trees found the rest of the Lophotrochozoa to consist of two monophyletic groups. The first group comprises of the Gastrotricha (Mesodasys laticaudatus), and the Platyhelminthes (flatworms). This relationship is consistent with recent studies (Dunn et al., 2008; Edgecombe et al., 2011; Laumer et al., 2015; Struck et al., 2014). Because of their primitive nature, with characteristics such as having no body cavity, no respiratory organs, and having only a single opening for both the intake of nutrients and excretion of waste, they were originally thought to be amongst the more primitive Bilateria, until molecular studies on 18S rDNA sequence data was carried out, placing them within the protostomes (Baguñà and Riutort, 2004). Authors now divide the Platyhelminthes into the Catenulida, with currently no known synapomorphies, and the Rhabditophora, which has uniting characteristics such as the presence of lamellated rhabdites, a common structure of the epidermis (Egger et al., 2015; Laumer et al., 2015). Our ML and Bayesian trees corroborated this, and found the Catenulida (Catenulida sp.) to be sister to Rhabditophora (Macrostomum lignano, Echinoplana celerrima, Microdalyellia schmidti, Monocelis, Schmidtea mediterranea).

Within the Rhabditophora, the most basal branches are those of the Macrostomorpha (Macrostomum lignano), followed by the Polycladida (Echinoplana celerrima), also in agreement with recent studies (Egger et al., 2015; Laumer et al., 2015). We see a disagreement between the ML and Bayesian tree topologies regarding the rest of the Rhabditophora. The ML tree inferred the Proseriata (Monocelis sp.) to be more basal than both the Rhabdocoela (Microdalyellia schmidti) and the Acentrosomata (Schmidtea mediterranea). This is in disagreement with recent analyses (Egger et al., 2015; Laumer et al., 2015), which places the Rhabdocoela as the most basal, followed by Proseriata and then Acentrosomata. The tree found through Bayesian inference agrees with the pedlishub phylogenies, however. As with the placement of the Rotifera within the Ecdysozoans under the ML analysis, this is possibly due to the ML inference method being more susceptible to Long Branch Attraction artefacts than the Bayesian, leading the long branched Rhabdocoela to be attracted to the long branched Acentrosomata in this framework.

The second monophyletic group found within the rest of the Lophotrochozoa contains the Annelida (Lumbricus rubellus Helobdella robusta Capitella sp.), worms, the Mollusca (Biomphalaria glabrata, Lymnaea stagnalis, Lottia gigantea, Mytilus californianus, Sepia officinalis, Chaetopleura apiculata), the largest marine phylum, and Nemertea (Cerebratulus sp.), also known as ribbon worms or proboscis worms, to form the Trochozoa (Dunn et al., 2014). However, there is disagreement on the positioning of these clades within the group (Dunn et al., 2008; Laumer et al., 2015; Struck et al., 2014; Struck and Fisse, 2008). Both tree reconstruction methods find the Gastropoda (Lottia gigantea, Lymnaea stagnalis, Biomphalaria glabrata) to be sister to the Bivalvia (Mytilus californianus). Both methods also found the Annelida to be sister to (Mollusca + Nemertea), with high support (posterior probability of 1 and bootstrap of 100).

By contrast, trees obtained from other orthology pipelines had more unresolved nodes and/or more discrepancies with the literature (Figure 4; table 1).

**Table 2:** Summary of support for major clades in trees obtained using the different methods. P indicates presence of clsade in PhyloBayes trees (bold: posterior probability >=0.95). L indicates presence of clade in maximum likelihood tree (bold: branch support >=0.95)

| Hypothesis | OMA | OrthoMCL | HamSTR | BUSCO |
| --- | --- | --- | --- | --- |
| Monophyly of Lophotrochozoa ((Dunn et al., 2008; Kocot et al., 2017; Telford et al., 2015)) | **P**<br>- | **P**<br>- | **P**<br>- | **P**<br>- |
| Gastropoda sister to Bivalvia ((Kocot et al., 2011)) | **P**<br>**L** | -<br>L | -<br>- | -<br>- |
| Annelida to be sister to (Mollusca + Nemertea) (Egger et al., 2015) | **P**<br>**L** | **P**<br>**L** | -<br>**L** | **P**<br>**L** |
| Monophyly of Deuterostomes | **P**<br>L | -<br>L | **P**<br>**L** | **P**<br>**L** |
| Rotifera sister to rest of Lophotrochozoa (Laumer et al., 2015) | **P**<br>- | -<br>- | **P**<br>- | **P**<br>- |
| Catenulida sister to Rhabditophora (Egger et al., 2015) | **P**<br>**L** | **P**<br>L | **P**<br>**L** | **P**<br>**L** |
| Monophyly of Annelida | **P**<br>**L** | **P**<br>**L** | **P**<br>**L** | **P**<br>**L** |
| Sister Clade of Gastrotricha to Platyhelminthes (Egger et al., 2015; Laumer et al., 2015) | **P**<br>L | **P**<br>- | **P**<br>**L** | **P**<br>- |
| **Total majority outcomes** | **8** (PhyloBayes)<br>**6** (RaxML) | **5** (PhyloBayes)<br>**5** (RaxML) | **6** (PhyloBayes)<br>**5** (RaxML) | **7** (PhyloBayes)<br>**4** (RaxML) |

**Fig 4:**
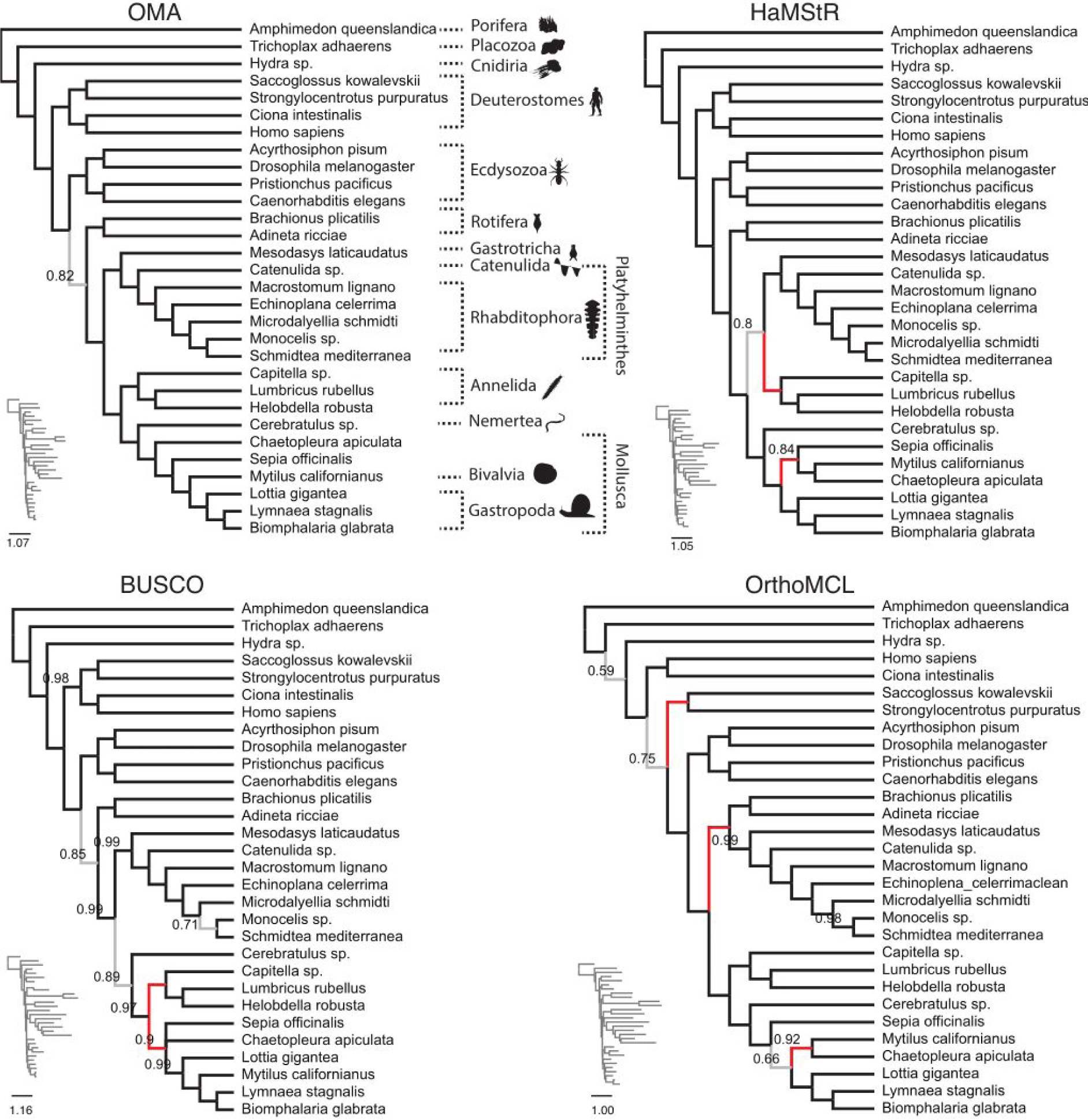
Comparison of trees obtained using PhyloBayes with the CAT-GTR-G4 model for different datasets. OMA tree is in congruence with published results (see main text). Branches which are at odds with the literature are in red; else they are displayed in grey (posterior probability < 0.95) or else in black. Only posterior probabilities below 1 are displayed.

The BUSCO Bayesian tree had slightly less support throughout than the OMA tree, although only had one branch with support of less than pp=0.80. The relationship between the Poreriata, Rhabdocoela and the Acentrosomata agrees with the OMA Bayesian tree, as does the relationship between the Gastrotricha and the Platyhelminthes. However, the BUSCO tree indicates Gastropoda to be paraphyletic with high support (pp=0.99), with Lottia gigantea to be more basal to the Bivalvia and the rest of the Gastropoda. This is in contrast to both the OMA tree and other studies (Dunn et al., 2008; Struck et al., 2014). The BUSCO tree found the Nemertea as sister to (Annelida + Mollusca), with a support value of pp=0.89. This is in disagreement with current consensus, and the OMA tree (Dunn et al., 2008; Laumer et al., 2015; Struck et al., 2014).

The HaMStR tree had high support throughout, but differed from the OMA tree. The HaMStR method placed the Sepia officinalis, Mytilus californianus and the Chaetopleura apiculata in a clade together, sister to the Gastropoda. This is in disagreement with (Kocot et al., 2011) and the OMA trees, which place the Polyplacophora (Chaetopleura apiculata) to be the most basal, followed by the Cephalopoda (Sepia officinalis), with the Bivalvia sister to the Gastropoda. The Bayesian tree also fails to recover the Trochozoa, placing the Annelida with the (Platyhelminthes+Gastrotricha), as opposed to full support found in the OMA tree.

The OrthoMCL trees had the most issues, with the lowest support values. Deuterostomes, comprising of a well established relationship between the Chordates and the Ambulacraria (Philippe et al., 2011a), are paraphyletic in the Phylobayes tree, which places the Chordates (Ciona intestinalis, Homo sapiens) basal to the Ambulacraria (Strongylocentrotus purpuratus, Saccoglossus kowalevskii), with the latter sister to the Protostomes with pp=0.75. The Rotifera were incorrectly placed as sister to (Gastrotricha + Platyhelminthes) with full support. This is in disagreement with both the OMA tree and recent studies. The tree was able to correctly infer the (Mollusca + Nemertea) relationship with full support. Within the Mollusca, in contrast to the OMA tree, the Bayesian tree inferred the Sepia officinalis to be the most basal, with Chaetopleura apiculata and Mytilus californianus forming a clade sister to the rest of the Mollusca. However, this has low support with pp=0.66 for the Bayesian tree.

## Discussion and outlook

OMA standalone enables researchers to infer high-quality orthologs among genomes or transcriptomes, on public and in house data. It runs on a wide range of hardwares, from a single computer to large clusters with thousands of parallel processes.

On the Lophotrochozoa dataset, compared with other approaches, OMA yielded more orthologous information for phylogenetic species tree inference and resulted in better resolved trees which are more consistent with the existing literature.

OMA standalone was also successful used to analyse centipedes (Fernández et al., 2014), arachnids (Fernández and Giribet, 2015; Sharma et al., 2014), assassin flies (Dikow et al., 2017), scorpions (Sharma et al., 2015), spiders (Garrison et al., 2016), flatworms (Egger et al., 2015; Laumer et al., 2015), tapeworms (Tsai et al., 2013), or Archaea (Williams et al., 2017).

Beyond species tree inference, OMA can also be used to pinpoint the emergence of gene families in evolution, an approach that is sometimes referred to as phylostratigraphy (Domazet-Lošo et al., 2007). Conventional approaches work by considering all the genes annotated in a species of reference, and performing BLAST searches against increasingly distant sets of taxa. The point at which no homolog can be found is inferred to immediately precede the emergence of the gene. However, such an approach does not differentiate between orthologs and paralogs, and thus has a limited resolution in terms of subfamilies. Alternatively, it is possible to extract more fine-grained information from reconciled gene trees (Huerta-Cepas et al., 2014; e.g. Vilella et al., 2008), but this is computationally demanding and there is a lack of tools to perform such analyses on custom data.

By inferring high-quality hierarchical orthologous groups, OMA standalone provides a way to map gene emergence, gene duplication, and gene loss onto species phylogenies. For instance, OMA standalone has been used to contrast gene families that have expanded and contracted in the common ancestors of echolocating and non-echolocating bats. The emergence of echolocation coincides with a decrease in chemosensory genes, while secondary loss of echolocation coincides with an increase in chemosensory genes (Tsagkogeorga et al., 2017).

For neglected tropical diseases, which disproportionately affect poorer people, it can be challenging to develop new medicines. To accelerate drug development in such cases, drug repurposing has been suggested whereby an already existing and approved medicine, or a well researched lead, is used to combat neglected tropical diseases (Ekins et al., 2011). Closantel, a veterinary anthelmintic has, for instance, been suggested for treatment of the human disease river blindness, caused by the filarial nematode Onchocerca volvulus (Gloeckner et al., 2010). As a first-pass bioinformatic identification of drug targets in four newly sequenced tapeworm genomes, OMA standalone was used to identify orthologs of known human drug targets (Tsai et al., 2013): Human genes targeted by drugs were retrieved from various databases, and their orthologs in tapeworms were inferred using OMA standalone. To identify targets likely to be essential across animals, orthologs with mice and nematodes were also identified: if both mice and nematode orthologs had knock-out phenotypes, we inferred that the orthologous group was essential across animals. Together with other indicators, such as gene expression data, we were able to rank every gene in these largely unexplored genomes for their suitability as a drug target, and associate lead compounds to them. As drugs could exhibit off-target effects on paralogs, the analysis focused on orthologs, which tend to be functionally more conserved (e.g. Altenhoff et al., 2012). The importance of investigating orthologs was illustrated by the drug Praziquantel, which is efficient against adult tapeworms, but not against the more dangerous larval form (Nogi et al., 2009). Praziquantel targets one particular voltage-gated calcium channel subunit. Using OMA standalone, we could identify the precise subunit ortholog in tapeworms and show that it is not expressed in the larval form—thereby providing a plausible explanation for the drug’s low efficacy.

To conclude, orthology inference is a key step in integrating biological knowledge across multiple species. OMA standalone is a versatile orthology inference software with a proven track record. The software has been continuously improved and maintained over the past five years, undergoing 2 major and 25 minor (bug fixing) releases. We intend to keep developing and maintaining it. For support enquiries or bug reporting, we encourage users to use the biostars.org forum using the keyword “oma”.

## Material and Methods

### Large-scale species phylogenetic reconstruction: Lophotrochozoa

We used transcriptome from seven Lophotrochozoa species published in (Egger et al., 2015): Mesodasys laticaudatus (Gastrotricha), Catenulida sp., Macrostomum ligano, Echinoplana celerrima, Microdalyellia schmidti, Monocelis sp. (Platyhelminthes) and Cerebratulus sp. (Nemertea). In addition, 12 sets of genomic and transcriptomic protein predictions from Saccoglossus kowalevskii, Brachionus plicatilis, Adineta ricciae, Schmidtea mediterranea, Lumbricus rubellus, Chaetopleura apiculata, Sepia officinalis, Mytilus californianus, Biomphalaria glabrata, Lymnaea stagnalis, Hydra magnipapillata and Amphimedon queenslandica were downloaded from the NCBI refseq repository (ftp.ncbi.nlm.nih.gov/refseq/). Redundant sequences with higher than 97% identity were removed by clustering with CD-HIT (Fu et al., 2012). Additionally, 11 precomputed proteomes for Homo sapiens, Strongylocentrotus purpuratus, Ciona intestinalis, Trichoplax adhaerens, Pristionchus pacificus, Caenorhabditis elegans, Drosophila melanogaster, Acyrthosiphon pisum, Capitella sp., Helobdella robusta and Lottia gigantea were downloaded from the OMA database website. The combined set of 30 non-redundant proteins sets contained 19 lophotrochozoans, four deuterostomes, four ecdysozoans, and proteomes from three non-bilaterian animals.

Quality assessment of sequencing reads was carried out with FastQC (Andrews and Others, 2010). Subsequent to this, it was determined, using PRINSEQ lite (Schmieder and Edwards, 2011), that the first 12 nucleotides should be trimmed off the 100bp reads. The assembly of the trimmed paired reads was done using Trinity v20130225 (Haas et al., 2013), with the flag ‘--min_kmer_cov 2’, with default parameters.

In order to detect the presence of cross contaminations between the various libraries run on the same flow cell, we used the CroCo package (Simion et al., 2018). This identified any assembled transcripts with fewer than four read matches, which were subsequently discarded. Furthermore, this also discarded all transcripts in which the number of reads, from the intended species matching the transcript, was not at least five times greater than the number of matches to the transcript, from reads from any of the other potentially contaminating species.

For peptide predictions, all ORFs greater than 100aa were retained. For all peptide datasets, cd-hit was used to reduce redundancy by clustering sequences with a global sequence identity of greater than 95%.

For the HaMStR analysis, putative orthologs were determined for each species using HaMStR v13.2.6 (Ebersberger et al., 2009) using the Lophotrochozoa core ortholog set.

Orthologous groups were inferred by running BUSCO v1.22 (Simão et al., 2015) on the Metazoa dataset found at (https://busco.ezlab.org/v1/). We created orthologous groups made up of the protein sequences which BUSCO deemed to have had complete matches with their own highly conserved genes. At most one species containing multiple sequences was allowed per group. There was only a single occurrence of a group containing multiple sequences from a single species. In this case, we retained only the longest sequence.

The set of 30 proteomes were first filtered to remove low quality protein sequences using the OrthoMCL script “orthomclFilterFasta” (Chen et al., 2006). Low quality sequences were defined to be sequences that were shorter than 10 amino acids, contained more than 20% stop codons, and contained more than 20% non-standard amino acids. An all versus all NCBI BLAST was then used with default parameters, in order to find the similarity score between sequences. Matches with an E-value < 10^-6^ were retained. Orthologs, in-paralogs and co-orthologs were then identified using the OrthoMCL script “OrthomclPairs” before clustering using MCL. An MCL inflation parameter of 2.2 was used in order to identify clusters. Each group was required to have at most one species containing multiple sequences. When more than one sequence from a single species was present, the longest sequence was selected to remain in the group, with the others removed.

Each orthologous group which contained a minimum of 15 protein sequences, of the 30 total, representing unique species were aligned using MUSCLE (Edgar, 2004), using default parameters. All spurious sequences, and poorly aligned regions of the multiple sequence alignments, were then removed using trimAl (Capella-Gutiérrez et al., 2009), using the -automated1 flag. Supermatrices were then constructed by concatenating all of the remaining alignments, with missing sequences treated as gaps. The final alignment was subsequently reduced to only contain sites in which more than 60% were occupied by amino acids.

Species trees were constructed using an LG+I model with 100 bootstrap replicates and a CAT+GTR+G4 model, with RAxMLv8.2.4 and PhyloBayes MPI v1.5a respectively. Convergence information is provided in Table 2.

**Table 2:** Convergence of the PhyloBayes runs

| Method | Num Cycles | MaxDiff | MeanDiff |
| --- | --- | --- | --- |
| OMA | 7,080 | 0.297691 | 0.00522266 |
| HaMStR | 2,731 | 0.297693 | 0.00972485 |
| BUSCO | 47,281 | 0.0957351 | 0.00435825 |
| OrthoMCL | 3,190 | 0.104071 | 0.0548237 |

## Acknowledgements

Computations were performed on the University College London Computer Science cluster and at the Vital-IT Center for high-performance computing of the SIB Swiss Institute of Bioinformatics. J.L is funded by EPSRC Centre for Doctoral Training studentship at UCL CoMPLEX (EP/F500351/1). MT acknowledges support by a Biotechnology and Biological Sciences Research Council grant (BB/H006966/1) and the European Research Council (ERC-2012-AdG 322790). C.D. acknowledges support by Swiss National Science Foundation grant 150654, UK BBSRC grant, and the Swiss State Secretariat for Education, Research and Innovation (SERI).

---

**Fig. S1:**
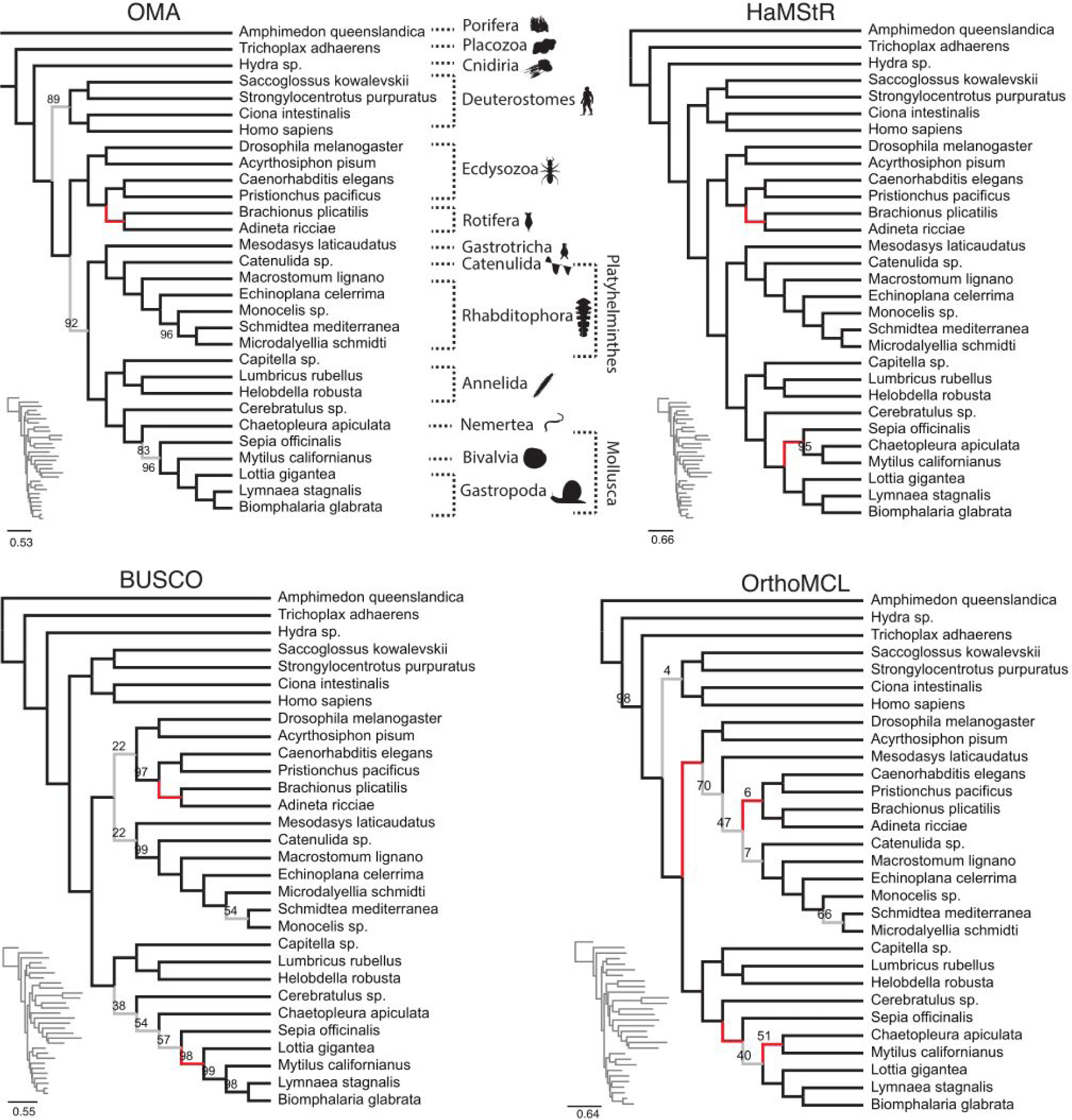
Comparison of trees obtained using RAxML with the LG+I model for different datasets. OMA tree is in congruence with published results (see main text). Branches which are at odds with the literature are in red; else they are displayed in grey (posterior probability < 0.95) or else in black. Only posterior probabilities below 1 are displayed.

## References

Afrasiabi C, Samad B, Dineen D, Meacham C, Sjölander K. 2013. The PhyloFacts FAT-CAT web server: ortholog identification and function prediction using fast approximate tree classification. Nucleic Acids Res 41:W242–8.

Altenhoff AM, Boeckmann B, Capella-Gutierrez S, Dalquen DA, DeLuca T, Forslund K, Huerta-Cepas J, Linard B, Pereira C, Pryszcz LP, Schreiber F, da Silva AS, Szklarczyk D, Train C-M, Bork P, Lecompte O, von Mering C, Xenarios I, Sjölander K, Jensen LJ, Martin MJ, Muffato M, Quest for Orthologs consortium, Gabaldón T, Lewis SE, Thomas PD, Sonnhammer E, Dessimoz C. 2016. Standardized benchmarking in the quest for orthologs. Nat Methods 13:425–430.

Altenhoff AM, Dessimoz C. 2012. Inferring Orthology and Paralogy In: Anisimova M, editor. Evolutionary Genomics, Methods in Molecular Biology. Humana Press. pp. 259–279.

Altenhoff AM, Dessimoz C. 2009. Phylogenetic and functional assessment of orthologs inference projects and methods. PLoS Comput Biol 5:e1000262.

Altenhoff AM, Gil M, Gonnet GH, Dessimoz C. 2013. Inferring hierarchical orthologous groups from orthologous gene pairs. PLoS One 8:e53786.

Altenhoff AM, Glover NM, Train C-M, Kaleb K, Warwick Vesztrocy A, Dylus D, de Farias TM, Zile K, Stevenson C, Long J, Redestig H, Gonnet GH, Dessimoz C. 2017. The OMA orthology database in 2018: retrieving evolutionary relationships among all domains of life through richer web and programmatic interfaces. Nucleic Acids Res. doi:10.1093/nar/gkx1019

Altenhoff AM, Škunca N, Glover N, Train C-M, Sueki A, Piližota I, Gori K, Tomiczek B, Müller S, Redestig H, Gonnet GH, Dessimoz C. 2015. The OMA orthology database in 2015: function predictions, better plant support, synteny view and other improvements. Nucleic Acids Res 43:D240–9.

Altenhoff AM, Studer RA, Robinson-Rechavi M, Dessimoz C. 2012. Resolving the ortholog conjecture: orthologs tend to be weakly, but significantly, more similar in function than paralogs. PLoS Comput Biol 8:e1002514.

Andrews S, Others. 2010. FastQC: a quality control tool for high throughput sequence data.

Baguñà J, Riutort M. 2004. Molecular phylogeny of the Platyhelminthes. Can J Zool 82:168–193.

Boeckmann B, Robinson-Rechavi M, Xenarios I, Dessimoz C. 2011. Conceptual framework and pilot study to benchmark phylogenomic databases based on reference gene trees. Brief Bioinform 12:423–435.

Capella-Gutiérrez S, Silla-Martínez JM, Gabaldón T. 2009. trimAl: a tool for automated alignment trimming in large-scale phylogenetic analyses. Bioinformatics 25:1972–1973.

Carrigan MA, Uryasev O, Davis RP, Zhai L, Hurley TD, Benner SA. 2012. The natural history of class I primate alcohol dehydrogenases includes gene duplication, gene loss, and gene conversion. PLoS One 7:e41175.

Chen F, Mackey AJ, Stoeckert CJ, Roos DS. 2006. OrthoMCL-DB: querying a comprehensive multi-species collection of ortholog groups. Nucleic Acids Res 34:D363–8.

Dalquen D a., Dessimoz C. 2013. Bidirectional Best Hits Miss Many Orthologs in Duplication-Rich Clades such as Plants and Animals. Genome Biol Evol 5:1800–1806.

Dessimoz C, Boeckmann B, Roth ACJ, Gonnet GH. 2006. Detecting non-orthology in the COGs database and other approaches grouping orthologs using genome-specific best hits. Nucleic Acids Res 34:3309–3316.

Dessimoz C, Cannarozzi G, Gil M, Margadant D, Roth A, Schneider A, Gonnet G. 2005. OMA, A Comprehensive, Automated Project for the Identification of Orthologs from Complete Genome Data: Introduction and First Achievements In: McLysaght A, Huson DH, editors. RECOMB 2005 Workshop on Comparative Genomics. Springer-Verlag. pp. 61–72.

Dessimoz C, Gabaldón T, Roos DS, Sonnhammer ELL, Herrero J, Quest for Orthologs Consortium. 2012. Toward community standards in the quest for orthologs. Bioinformatics 28:900–904.

Dikow RB, Frandsen PB, Turcatel M, Dikow T. 2017. Genomic and transcriptomic resources for assassin flies including the complete genome sequence of Proctacanthus coquilletti (Insecta: Diptera: Asilidae) and 16 representative transcriptomes. PeerJ 5:e2951.

Domazet-Lošo T, Brajković J, Tautz D. 2007. A phylostratigraphy approach to uncover the genomic history of major adaptations in metazoan lineages. Trends Genet 23:533–539.

Dunn CW, Giribet G, Edgecombe GD, Hejnol A. 2014. Animal Phylogeny and Its Evolutionary Implications*. Annu Rev Ecol Evol Syst 45:371–395.

Dunn CW, Hejnol A, Matus DQ, Pang K, Browne WE, Smith SA, Seaver E, Rouse GW, Obst M, Edgecombe GD, Sørensen MV, Haddock SHD, Schmidt-Rhaesa A, Okusu A, Kristensen RM, Wheeler WC, Martindale MQ, Giribet G. 2008. Broad phylogenomic sampling improves resolution of the animal tree of life. Nature 452:745–749.

Ebersberger I, Strauss S, von Haeseler A. 2009. HaMStR: profile hidden markov model based search for orthologs in ESTs. BMC Evol Biol 9:157.

Edgar RC. 2004. MUSCLE: multiple sequence alignment with high accuracy and high throughput. Nucleic Acids Res 32:1792–1797.

Edgecombe GD, Giribet G, Dunn CW, Hejnol A, Kristensen RM, Neves RC, Rouse GW, Worsaae K, Sørensen MV. 2011. Higher-level metazoan relationships: recent progress and remaining questions. Org Divers Evol 11:151–172.

Egger B, Lapraz F, Tomiczek B, Müller S, Dessimoz C, Girstmair J, Škunca N, Rawlinson KA, Cameron CB, Beli E, Todaro MA, Gammoudi M, Noreña C, Telford MJ. 2015. A Transcriptomic-Phylogenomic Analysis of the Evolutionary Relationships of Flatworms. Curr Biol 0. doi:10.1016/j.cub.2015.03.034

Ekins S, Williams AJ, Krasowski MD, Freundlich JS. 2011. In silico repositioning of approved drugs for rare and neglected diseases. Drug Discov Today 16:298–310.

Fernández R, Giribet G. 2015. Unnoticed in the tropics: phylogenomic resolution of the poorly known arachnid order Ricinulei (Arachnida). R Soc Open Sci 2:150065.

Fernández R, Laumer CE, Vahtera V, Libro S, Kaluziak S, Sharma PP, Pérez-Porro AR, Edgecombe GD, Giribet G. 2014. Evaluating topological conflict in centipede phylogeny using transcriptomic data sets. Mol Biol Evol 31:1500–1513.

Forslund K, Pereira C, Capella-Gutierrez S, Sousa da Silva A, Altenhoff A, Huerta-Cepas J, Muffato M, Patricio M, Vandepoele K, Ebersberger I, Blake J, Fernández Breis JT, Quest for Orthologs Consortium, Boeckmann B, Gabaldón T, Sonnhammer E, Dessimoz C, Lewis S. 2017. Gearing up to handle the mosaic nature of life in the quest for orthologs. Bioinformatics. doi:10.1093/bioinformatics/btx542

Fu L, Niu B, Zhu Z, Wu S, Li W. 2012. CD-HIT: accelerated for clustering the next-generation sequencing data. Bioinformatics 28:3150–3152.

Gabaldón T, Koonin EV. 2013. Functional and evolutionary implications of gene orthology. Nat Rev Genet 14:360–366.

Garrison NL, Rodriguez J, Agnarsson I, Coddington JA, Griswold CE, Hamilton CA, Hedin M, Kocot KM, Ledford JM, Bond JE. 2016. Spider phylogenomics: untangling the Spider Tree of Life. PeerJ 4:e1719.

Giribet G, Edgecombe GD. 2017. Current Understanding of Ecdysozoa and its Internal Phylogenetic Relationships. Integr Comp Biol 57:455–466.

Gloeckner C, Garner AL, Mersha F, Oksov Y, Tricoche N, Eubanks LM, Lustigman S, Kaufmann GF, Janda KD. 2010. Repositioning of an existing drug for the neglected tropical disease Onchocerciasis. Proc Natl Acad Sci U S A 107:3424–3429.

Haas BJ, Papanicolaou A, Yassour M, Grabherr M, Blood PD, Bowden J, Couger MB, Eccles D, Li B, Lieber M, Macmanes MD, Ott M, Orvis J, Pochet N, Strozzi F, Weeks N, Westerman R, William T, Dewey CN, Henschel R, Leduc RD, Friedman N, Regev A. 2013. De novo transcript sequence reconstruction from RNA-seq using the Trinity platform for reference generation and analysis. Nat Protoc 8:1494–1512.

Hejnol A, Obst M, Stamatakis A, Ott M, Rouse GW, Edgecombe GD, Martinez P, Baguñà J, Bailly X, Jondelius U, Wiens M, Müller WEG, Seaver E, Wheeler WC, Martindale MQ, Giribet G, Dunn CW. 2009. Assessing the root of bilaterian animals with scalable phylogenomic methods. Proc Biol Sci 276:4261–4270.

Huerta-Cepas J, Capella-Gutiérrez S, Pryszcz LP, Marcet-Houben M, Gabaldón T. 2014. PhylomeDB v4: zooming into the plurality of evolutionary histories of a genome. Nucleic Acids Res 42:D897–902.

Huerta-Cepas J, Szklarczyk D, Forslund K, Cook H, Heller D, Walter MC, Rattei T, Mende DR, Sunagawa S, Kuhn M, Jensen LJ, von Mering C, Bork P. 2016. eggNOG 4.5: a hierarchical orthology framework with improved functional annotations for eukaryotic, prokaryotic and viral sequences. Nucleic Acids Res 44:D286–93.

Kocot KM. 2016. On 20 years of Lophotrochozoa. Org Divers Evol 16:329–343.

Kocot KM, Cannon JT, Todt C, Citarella MR, Kohn AB, Meyer A, Santos SR, Schander C, Moroz LL, Lieb B, Halanych KM. 2011. Phylogenomics reveals deep molluscan relationships. Nature 477:452–456.

Kocot KM, Struck TH, Merkel J, Waits DS, Todt C, Brannock PM, Weese DA, Cannon JT, Moroz LL, Lieb B, Halanych KM. 2017. Phylogenomics of Lophotrochozoa with Consideration of Systematic Error. Syst Biol 66:256–282.

Lartillot N, Rodrigue N, Stubbs D, Richer J. 2013. PhyloBayes MPI: phylogenetic reconstruction with infinite mixtures of profiles in a parallel environment. Syst Biol 62:611–615.

Laumer CE, Hejnol A, Giribet G. 2015. Nuclear genomic signals of the “microturbellarian” roots of platyhelminth evolutionary innovation. Elife 4. doi:10.7554/eLife.05503

Li L, Stoeckert CJ, Roos DS. 2003. OrthoMCL: identification of ortholog groups for eukaryotic genomes. Genome Res 13:2178–2189.

Linard B, Thompson JD, Poch O, Lecompte O. 2011. OrthoInspector: comprehensive orthology analysis and visual exploration. BMC Bioinformatics 12:11.

Mi H, Huang X, Muruganujan A, Tang H, Mills C, Kang D, Thomas PD. 2017. PANTHER version 11: expanded annotation data from Gene Ontology and Reactome pathways, and data analysis tool enhancements. Nucleic Acids Res 45:D183–D189.

Nogi T, Zhang D, Chan JD, Marchant JS. 2009. A novel biological activity of praziquantel requiring voltage-operated Ca2+ channel beta subunits: subversion of flatworm regenerative polarity. PLoS Negl Trop Dis 3:e464.

Pagani I, Liolios K, Jansson J, Chen I-MA, Smirnova T, Nosrat B, Markowitz VM, Kyrpides NC. 2012. The Genomes OnLine Database (GOLD) v.4: status of genomic and metagenomic projects and their associated metadata. Nucleic Acids Res 40:D571–9.

Pellegrini M, Marcotte EM, Thompson MJ, Eisenberg D, Yeates TO. 1999. Assigning protein functions by comparative genome analysis: protein phylogenetic profiles. Proc Natl Acad Sci U S A 96:4285–4288.

Philippe H, Brinkmann H, Copley RR, Moroz LL, Nakano H, Poustka AJ, Wallberg A, Peterson KJ, Telford MJ. 2011a. Acoelomorph flatworms are deuterostomes related to Xenoturbella. Nature 470:255–258.

Philippe H, Brinkmann H, Lavrov DV, Littlewood DTJ, Manuel M, Wörheide G, Baurain D. 2011b. Resolving difficult phylogenetic questions: why more sequences are not enough. PLoS Biol 9:e1000602.

Roth AC, Gonnet GH, Dessimoz C. 2008. Correction: Algorithm of OMA for large-scale orthology inference. BMC Bioinformatics 9:518.

Schmieder R, Edwards R. 2011. Quality control and preprocessing of metagenomic datasets. Bioinformatics 27:863–864.

Schmitt T, Messina DN, Schreiber F, Sonnhammer ELL. 2011. Letter to the editor: SeqXML and OrthoXML: standards for sequence and orthology information. Brief Bioinform 12:485–488.

Sharma PP, Fernandez R, Santillan GR, Monod L. 2015. Phylogenomic resolution of scorpions reveals discordance with morphological phylogenetic signalINTEGRATIVE AND COMPARATIVE BIOLOGY. OXFORD UNIV PRESS INC JOURNALS DEPT, 2001 EVANS RD, CARY, NC 27513 USA. pp. E165–E165.

Sharma PP, Kaluziak ST, Pérez-Porro AR, González VL, Hormiga G, Wheeler WC, Giribet G. 2014. Phylogenomic interrogation of arachnida reveals systemic conflicts in phylogenetic signal. Mol Biol Evol 31:2963–2984.

Simão FA, Waterhouse RM, Ioannidis P, Kriventseva EV, Zdobnov EM. 2015. BUSCO: assessing genome assembly and annotation completeness with single-copy orthologs. Bioinformatics 31:3210–3212.

Simion P, Belkhir K, François C, Veyssier J, Rink JC, Manuel M, Philippe H, Telford MJ. 2018. A software tool “CroCo” detects pervasive cross-species contamination in next generation sequencing data. BMC Biol 16:28.

Smith TF, Waterman MS. 1981. Identification of common molecular subsequences. J Mol Biol 147:195–197.

Sonnhammer ELL, Gabaldón T, Sousa da Silva AW, Martin M, Robinson-Rechavi M, Boeckmann B, Thomas PD, Dessimoz C, Quest for Orthologs consortium. 2014. Big data and other challenges in the quest for orthologs. Bioinformatics 30:2993–2998.

Sonnhammer ELL, Östlund G. 2015. InParanoid 8: orthology analysis between 273 proteomes, mostly eukaryotic. Nucleic Acids Res 43:D234–9.

Stamatakis A. 2014. RAxML version 8: a tool for phylogenetic analysis and post–analysis of large phylogenies. Bioinformatics 30:1312–1313.

Struck TH, Fisse F. 2008. Phylogenetic position of Nemertea derived from phylogenomic data. Mol Biol Evol 25:728–736.

Struck TH, Wey-Fabrizius AR, Golombek A, Hering L, Weigert A, Bleidorn C, Klebow S, Iakovenko N, Hausdorf B, Petersen M, Kück P, Herlyn H, Hankeln T. 2014. Platyzoan paraphyly based on phylogenomic data supports a noncoelomate ancestry of spiralia. Mol Biol Evol 31:1833–1849.

Szalkowski A, Ledergerber C, Krähenbühl P, Dessimoz C. 2008. SWPS3 - fast multi-threaded vectorized Smith-Waterman for IBM Cell/B.E. and x86/SSE2. BMC Res Notes 1:107.

Telford MJ, Budd GE, Philippe H. 2015. Phylogenomic Insights into Animal Evolution. Curr Biol 25:R876–87.

Train C-M, Glover NM, Gonnet GH, Altenhoff AM, Dessimoz C. 2017. Orthologous Matrix (OMA) algorithm 2.0: more robust to asymmetric evolutionary rates and more scalable hierarchical orthologous group inference. Bioinformatics 33:i75–i82.

Tsagkogeorga G, Müller S, Dessimoz C, Rossiter SJ. 2017. Comparative genomics reveals contraction in olfactory receptor genes in bats. Sci Rep 7:259.

Tsai IJ, Zarowiecki M, Holroyd N, Garciarrubio A, Sanchez-Flores A, Brooks KL, Tracey A, Bobes RJ, Fragoso G, Sciutto E, Aslett M, Beasley H, Bennett HM, Cai J, Camicia F, Clark R, Cucher M, De Silva N, Day T a., Deplazes P, Estrada K, Fernández C, Holland PWH, Hou J, Hu S, Huckvale T, Hung SS, Kamenetzky L, Keane J a., Kiss F, Koziol U, Lambert O, Liu K, Luo X, Luo Y, Macchiaroli N, Nichol S, Paps J, Parkinson J, Pouchkina-Stantcheva N, Riddiford N, Rosenzvit M, Salinas G, Wasmuth JD, Zamanian M, Zheng Y, Cai X, Soberón X, Olson PD, Laclette JP, Brehm K, Berriman M. 2013. The genomes of four tapeworm species reveal adaptations to parasitism. Nature 496:57–63.

Uchiyama I, Mihara M, Nishide H, Chiba H. 2012. MBGD update 2013: the microbial genome database for exploring the diversity of microbial world. Nucleic Acids Res 41:D631–5.

Vilella AJ, Severin J, Ureta-Vidal A, Heng L, Durbin R, Birney E. 2008. EnsemblCompara GeneTrees: Complete, duplication-aware phylogenetic trees in vertebrates. Genome Res 19:327–335.

Williams TA, Szöllősi GJ, Spang A, Foster PG, Heaps SE, Boussau B, Ettema TJG, Embley TM. 2017. Integrative modeling of gene and genome evolution roots the archaeal tree of life. Proceedings of the National Academy of Sciences. doi:10.1073/pnas.1618463114

Zdobnov EM, Tegenfeldt F, Kuznetsov D, Waterhouse RM, Simão FA, Ioannidis P, Seppey M, Loetscher A, Kriventseva EV. 2017. OrthoDB v9.1: cataloging evolutionary and functional annotations for animal, fungal, plant, archaeal, bacterial and viral orthologs. Nucleic Acids Res 45:D744–D749.

Zerbino DR, Achuthan P, Akanni W, Amode MR, Barrell D, Bhai J, Billis K, Cummins C, Gall A, Girón CG, Gil L, Gordon L, Haggerty L, Haskell E, Hourlier T, Izuogu OG, Janacek SH, Juettemann T, To JK, Laird MR, Lavidas I, Liu Z, Loveland JE, Maurel T, McLaren W, Moore B, Mudge J, Murphy DN, Newman V, Nuhn M, Ogeh D, Ong CK, Parker A, Patricio M, Riat HS, Schuilenburg H, Sheppard D, Sparrow H, Taylor K, Thormann A, Vullo A, Walts B, Zadissa A, Frankish A, Hunt SE, Kostadima M, Langridge N, Martin FJ, Muffato M, Perry E, Ruffier M, Staines DM, Trevanion SJ, Aken BL, Cunningham F, Yates A, Flicek P. 2018. Ensembl 2018. Nucleic Acids Res 46:D754–D761.

